# Investigation of the temporal and spatial dynamics of muscular action potentials through optically pumped magnetometers

**DOI:** 10.1101/2021.02.24.432771

**Authors:** Philip J. Broser, Justus Marquetand, Thomas Middelmann, Davide Sometti, Christoph Braun

## Abstract

**Aim:** This study aims to simultaneously record the magnetic and electric components of the propagating muscular action potential.

**Method:** A single-subject study of the monosynaptic stretch reflex of the musculus rectus femoris was performed; the magnetic field generated by the muscular activity was recorded in all three spatial directions by five optically pumped magnetometers. In addition, the electric field was recorded by four invasive fine-wire needle electrodes. The magnetic and electric fields were compared, and modelling and simulations were performed to compare the magnetic field vectors with the underlying muscular anatomy of the rectus femoris muscle.

**Results:** The magnetomyography (MMG) signal can reliably be recorded following the stimulation of the monosynaptic stretch reflex. The MMG signal shows several phases of activity inside the muscle, the first of which is the propagating muscular action potential. As predicted by the finite wire model, the magnetic field vectors of the propagating muscular action potential are generated by the current flowing longitudinal to the muscle fiber. Based on the magnetic field vectors, it was possible to reconstruct the pinnation angle in the muscle. The later magnetic components are linked to the activated contractile apparatus.

**Interpretation:** MMG allows to analyze the muscle physiology from the propagating muscular action potential to the initiation of the contractile apparatus. At the same time this methods reveal information about muscle fiber direction and extend. With the development of high-resolution magnetic cameras, it will be possible to image the function and structure of any skeletal muscle with high precision. This method could be used in clinical medicine but also in sports and training science.

**What this paper adds:**

- A robust technique for triggering a muscular action potential that can be recorded by MMG and needle EMG simultaneously
- The correlation of the MMG signal with the needle EMG signal
- A method for detecting the direction of the propagating muscular action potential
- A method for correlating the magnetic field vectors with the pinnation angle of the examined muscle

## 1 Introduction

The smooth and controlled movements of the skeletal muscles require the well-controlled activation of the contractile apparatus along the whole length of the contracting muscle (Ghezzi, 1991). Here, the precise initiation of the molecular muscular contractile apparatus critically depends on the timing and spread of the excitation along the muscular fiber by the muscular action potential (Martonosi, 2000). After activation of the postsynaptic neuromuscular endplate by the motoneuron, the signal to initiate the contractile apparatus spreads along the muscular fiber through the muscular action potential (Henneberg, 1997). The action potential not only spreads in the longitudinal direction but also along the T-tubuli in the radial direction. The depolarization of the T-tubuli membrane leads to an opening of the ryanodine receptors and thus initiates a calcium influx in the cytoplasm from outside the cell as well as from the sarcoplasmic reticulum. The increase in the calcium concentration finally activates the muscle contraction (Martonosi, 2000).

The muscular action potential generates a measurable electric and magnetic field, and both fields disclose valuable physiological information (Farina, 2001; Broser, 2020). The electric field is, for instance, used in motor nerve conduction studies to determine the exact time the muscle is activated by the electrically stimulated motor nerve. Given that the signal is generated by the summation of the electric field of many neuro muscular units (Broser, 2020), the signal is called the compound muscle action potential (CMAP). When examining the precise timing of the muscular action potential of a distinct set of neuro muscular units, needle EMG recording is currently the technique of choice (Moritani, 2004).

However, this technique is painful and only allows recording at one or a few locations in the muscle. However, only recently it has been shown that recording the magnetic field of the muscular action potential allows a precise non-invasive analysis of the temporal and spatial dynamics (Broser, 2020). Further, the magnetic field can theoretically be linked to the current flowing inside the muscle fiber and thus reveal the spread of excitation during the propagation of the action potential.

In the pioneering study by Broser (2020), the magnetic field generated by the muscular action potential propagating along the muscle fibers of an intrinsic foot muscle was studied. In order to obtain highly synchronized neuromuscular units, the supplying nerve was electrically stimulated. This study showed that the muscular action potential can be recorded with a high signal-to-noise ratio. It further demonstrated that the position of the neuromuscular endplate can be localized by recording the magnetic field vectors.

So far the golden standard for the study of the muscular action potential is invasive needle myography. Optically pumped magnetometers are a promising alternative. How comparable needle EMG and OPM recording are needs to be evaluated. This is especially important when considering that the magnetic field reflects more the current than transmembrane voltage (as concentric needle EMG does).

Moreover, previous studies on MMG (Broser, 2018; Broser, 2020) relied on an artificial stimulus by electrically stimulating the supplying nerve rather than using a physiological paradigm involving intentional muscle activations or eliciting a monosynaptic reflex response. In addition, so far, only small intrinsic hand or foot muscles have been measured, with large and anatomically more complex muscles remaining to be studied. Therefore, we developed an experimental protocol and designed an experimental setup to record the magnetic components of the propagating muscle action potential (magnetomyogram, MMG) and, simultaneously, the monopolar fine-wire electromyogram (EMG) after monosynaptic reflex stimulation.

With this study we aimed to explore the full potential of the MMG technique. Especially to investigate if its possible to analyze both, the physiology of the muscular action potential and also the fiber structure of the muscle under investigation.

For our analysis, we therefore selected the rectus femoris muscle. The rectus femoris is a longitudinal fusiform muscle (Ward, 2008) that partially flexes the hip and extends the leg at the knee (see Figure 1A). The muscle starts at two locations: the anterior inferior iliac spine and the upper rim of the acetabulum. The distal tendon of the muscle is part of the M. quadriceps tendon, which becomes the patella ligament and which is connected to the tuberositas tibiae. The superficial fibers are arranged in a three-dimensional bipenniform manner (Ward, 2009; Blemker, 2006). This complex anatomical situation is outlined in Figure 1B and Figure 2 Panel A. The proximal fibers are angulated with an angle to the longitudinal axis of the muscle of about 14 ± 4 degrees (see Figure 2A). The distal fibers run roughly parallel to the longitudinal axis. In addition to the angulation in the coronal plane, the fibers are also angulated in the sagittal plane (see Figure 2 Panel B).

**Figure 1.**
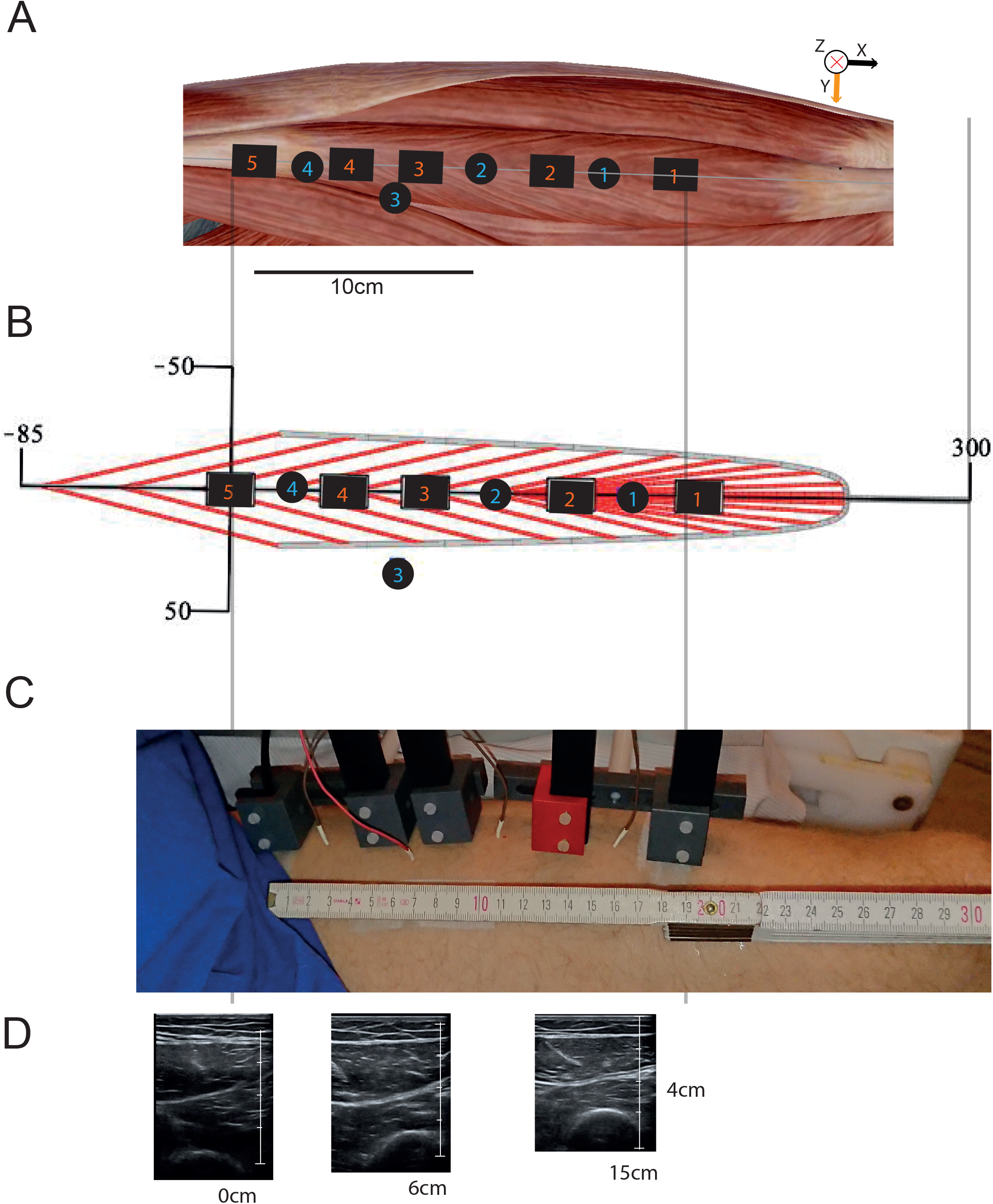
This figure shows the setup of the experiment and the relevant anatomical details. A: Macroscopic anatomy of the rectus femoris muscle. The black squares with orange numbers show the positions of the OPM sensor in relation to the muscle. The black circles with blue numbers show the positions of the fine-wire EMG needles. B: The rectus femoris muscle has a three-dimensional bipenniform structure. This structure is shown in this model, which will be used throughout the manuscript. C: Real-world image of the experiment setup. D: US sonographic images of the muscle at the depicted locations.

**Figure 2.**
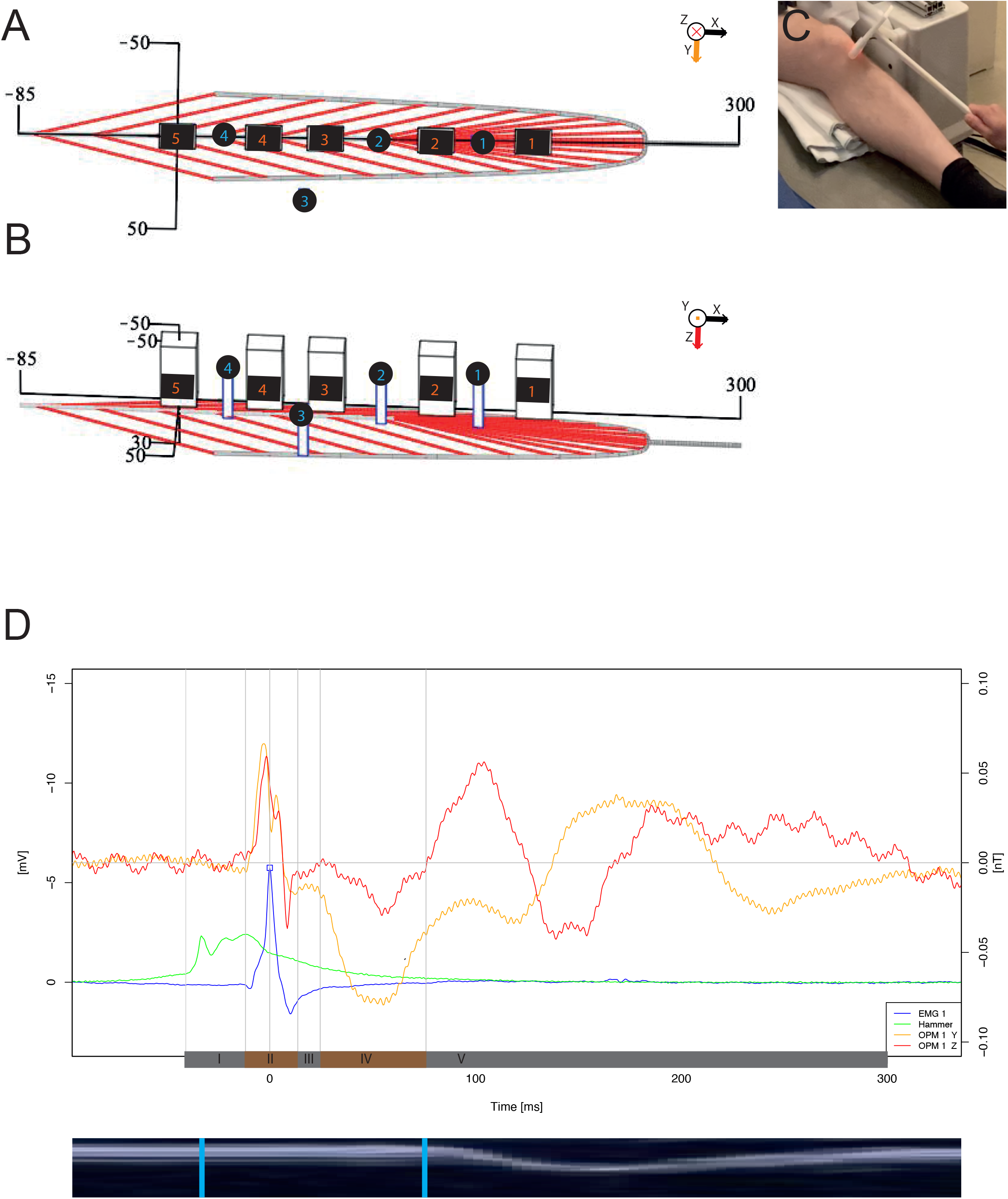
A, B: Modell of the rectus femoris. The model highlights the three-dimensional bipenniform structure of the muscle. The proximal fibers are angulated about 20° to the longitudinal extend; the distal fibers run almost parallel. C: Image showing the left leg of the subject resting on a pillow. In addition, the paramagnetic tendon hammer is shown in the picture hitting the patella tendon. D: Plot showing the signal traces of EMG 1 (blue), Hammer (green) and OPM 1 in the y- and z-direction. The identified distinct time intervals are shown at the bottom of the x axis labeled with roman numbers. At the bottom of the image, an M mode ultrasound image, which was temporally co-registered with the OPM data, is shown.

In this study we combined the OPM measurement of the magnetic field with fine-wire needle EMG electrodes (MedCat 0.4 × 10 mm Platin-Iridium needle) in order to measure the electric field at the same time. The muscle was activated by triggering the monosynaptic patellar tendon stretch reflex (Gürbüz, 2015) using a custom-made paramagnetic tendon hammer.

## 2 METHOD

### 2.1 Principal setup

A single-subject, experimental study was conducted at the MEG-Center of the University of Tübingen, Germany in October 2020. Experiments were performed according to the standards by the World Medical Association (World Medical Association, 2001). The subject of this study was an author of this publication, and he gave his consent for his data to published. The experiment was designed specifically to record the magnetic activity of the left rectus femoris muscle (Figure 1 Panel A) after triggering the monosynaptic patellar tendon stretch reflex (Figure 2 Panel B). Prior to the experiment, the muscle was imaged via high resolution muscle ultrasound (see Figure 1 Panel C) in order to determine the longitudinal axis of the muscle. To estimate the direction of the muscular fibers, a model based on the ideas in Blemker (2005) was developed and fitted to the subject’s body size using muscular ultrasound images. The electric activity of the muscle was recorded by four monopolar EMG needle electrodes (MedCat 0.4 × 10 mm Platin-Iridium needles), and the magnetic field was measured by five OPM sensors (Figure 1 Panel A and Panel C).

First, muscular ultrasound imaging (Mindray TE7, 14Mhz-linear probe, see Figure 1D) was used to measure the size and longitudinal length of the left rectus femoris muscle. The parameters of the muscular fiber model were adapted accordingly (see Figure 1B). After ultrasound imaging, the subject sat down on a comfortable chair inside a magnetic shielded room (Ak3b, VAC Vacuumschmelze, Hanau, Germany). The left leg of the subject was supported by a pillow (Figure 2C) so that the rectus femoris was completely relaxed and the patella tendon could be easily reached by an examinator using a tendon hammer (Figure 2 C). The hammer was equipped with fiber optics in order to produce a trigger signal at the time point the hammer made contact with the participant’s skin. To produce the trigger, one of the two fibers built into the hammer provided an amplitude modulated light beam, and the other measured the reflection of the beam (Keyence Digital Fiber Sensor FS-N10). Approaching the participant’s skin with the hammer caused an increase in the reflected intensity until the skin was touched. At that instant, the reflected intensity dropped sharply.

Four monopolar^1^ fine-wire EMG needles (MedCat 0,4 × 10 mm Platin-Iridium needle) were placed inside or near the rectus femoris muscle (Figure 1B and C).

If possible, the needle electrodes were placed in a distal to proximal order along a line parallel to the fiber direction estimated via ultrasound imaging (see Figure 1A and B). In addition, a surface reference electrode was placed on the lateral skin at the level of the left knee, and a ground electrode was placed on the skin of the right shoulder.

Five optically pumped magnetometers (four OPM [No. 1 to 4], QZFM-gen-1.5 and one OPM [No. 5], Generation QZFM-gen-2) (QuSpin Inc., Louisville, CO, USA) were placed in a distal to proximal order along a line (see Figure 1A and B) parallel to the longitudinal extend of the muscle. The magnetometers were placed about 2–3 mm above the skin surface and were based on an optically detected zero-field resonance in hot rubidium vapor, which was contained in a vapor cell measuring 3 × 3 × 3 mm^3^. The center of the cell had a distance of 6.2 mm to the exterior of the housing (6.5 mm in the case of OPM No. 5), which measured (Generation 1.0) 13 × 19 × 85 mm^3^ (12.4 × 16.6 × 24.4 mm^3^ in the case of OPM No. 5). The small size of the OPM sensors allows for easy handling and their flexible adaptation to specific geometrical situations (Boto et al., 2017, Osborne et al., 2018, Sander et al. 2020). OPMs are capable of measuring two components of the magnetic field vector: the y- and z-direction. They measure with a magnetic field sensitivity in the order of 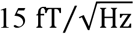 in a bandwidth of 3–135 Hz and a dynamic range of a few nanoteslas. To adapt to a non-zero magnetic background field, the sensors are equipped with internal compensation coils that can cancel magnetic background fields of up to 200 nT in the sensing vapor cell.

Our set of OPM sensors could simultaneously record the magnetic field in two orthogonal directions (y and z). In order to also record the x-direction, we repeated the measurements with the sensors turned by 90 degrees around the z-direction.

### 2.2 Stimulation

In order to record temporal synchronized muscular action potentials, the muscle was indirectly stimulated by eliciting the monosynaptic patellar stretch reflex (Gürbüz, 2015). To this end, a custom-made non-magnetic tendon hammer made of plastic was used (Figure 2C). The head of the hammer hosted two optical fibers. When the hammer was in close proximity to the skin, the light emitted from one of the fibers would be reflected by the skin back into the other fiber. Both fibers were tunneled out of the of the magnetic shielded room and connected to a light-emitting diode and a photo transistor. The voltage of the photo detector was amplified and fed into one analog-digital-converter (ADC) channel of the MEG recording system. Just a few millimeters before the hammer hit the skin, the amount of reflected light peaks—and, thus, the specific time points of the stimulation—could be determined (see Figure 2D). Several series of stimulations were recorded. First, a series was measured without boosting the reflex. Then, the reflex was enhanced by asking the subject to clench their teeth or by asking the subject to perform the Jendrassik maneuver. To grossly estimate the muscular anatomy during the contraction, a motion-mode (M-mode) ultrasound examination was done after the experiment. The M mode image was registered to the OPM and EMG data (see Figure 2D).

### 2.3 Recording

The data was recorded with the data acquisition function of the MEG system (CTF Omega 275, Coquitlam, BC, Canada) installed in the magnetically shielded rom. The analogue signals (OPMs, EMG, and Hammer) were digitized with a sampling rate of 2343.8 samples per second. A low-pass filter of 100 Hz was applied to the analogue signal of the OPMs. Four of the EEG channels included in the CFT MEG system were used to record the signals from the fine-wire needle EMGs. The EMG signals were band-pass filtered with a Butterworth filter with an edge frequency set at 5 Hz (high pass) and 800 Hz (low pass). The OPM system has an intrinsic delay between the measurement and analogue output; this delay was measured to be 3 ms. The data was post-hoc calibrated so that the magnetic field, electric potential, and signal of the hammer were in temporal synchrony.

### 2.4 Data processing and simulations

As described above, the data was recorded with the CTF System. The data was exported as continuous ASCII datafiles and imported to R (R Core Team, Vienna, Austria) for further processing. The signal from the hammer was used to separate the data into distinct trials (see Figure 2 D). In order to compare different trials with each other, the trial data was calibrated using the peak of the monopolar signal from EMG 1 as a reference (t = 0 ms, see Figure 2D). Apart from the offset calibration, no further processing was applied.

### 2.5 Modelling of the propagating muscular action potential

In addition to its temporal characteristics, the magnetic signal contains valuable geometric information. In order to link the obtained time-dependent magnetic fields with the underlying structural physiology, the finite wire model described in Broser (2020) was here applied, and parameter fitting was performed. The calculation was performed using Maple™ (Waterloo, ON, Canada). The finite wire model is based on the assumption that the muscular action potential propagates linearly along the muscle fiber. Therefore, the position of the action potential is parametrized by:

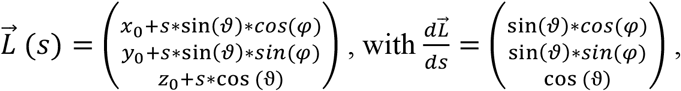

with L denoting the position of the muscular AP in space, s parameterizing the longitudinal extend, and the theta and phi angles defining the orientation of the fiber direction in polar coordinates. The model assumes—without loss of generality—that the recording sensor is located at position (0,0,0), with its z-direction pointing downwards.

The membrane potential orthogonal and longitudinal to the fiber orientation is modelled according to Rosenfalck (1969):

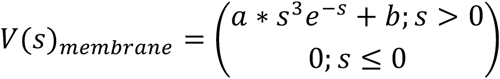

with 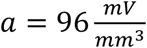 and *b* = −90 mV. The magnetic field then calculates to

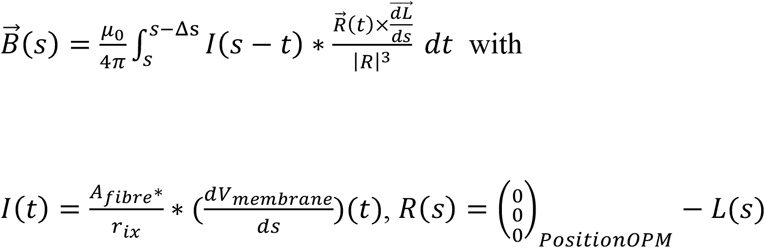

with *r_ix_* = 173 Ω ∗ *cm* = 17.3 Ω ∗ *mm* (Henneberg and Roberge, 1997), Δ*s* = 15 mm. A detailed description of the model can be found in Broser (2020).

## 3 Results

The subject had an upper-leg length of 30 cm (patella to groin). The upper aponeurosis of the M. rectus femoris muscle was 8 mm below the skin (see Figure 2D). After hitting the patellar tendon with the reflex hammer, a contraction of the rectus femoris muscle could be reliably triggered. Figure 2D shows the recorded signal of the hammer, EMG electrode 1, and the magneto-myogram (MMG) signal recorded with OPM 1 for the z- and y-direction in a time window of 400 ms around the time of the stimulation. In order to support the clarity of the manuscript, throughout the publication, the time window of the plots has been defined so that the highest negative peak in EMG electrode 1 for each trial defines the time point 0 ms. About 20 ms after the peak of the hammer signal, a small positive deflection followed by a large negative deflection can be detected in EMG channel 1. Simultaneously, a biphasic signal is recorded from OPM device No. 1 in the z- and y-direction. The biphasic signal is followed by a short phase of electric and magnetic silence lasting about 10 ms. Then, a large magnetic component in the y-direction and later in the z-direction can be distinguished. Table I lists the different time intervals that can be distinguished in the signals:

**Table I:** Time intervals of the MMG Signal

| Name | Time | Electric (EMG I monopolar) | Magnetic (OPM I Z and Y direction) | Description |
| --- | --- | --- | --- | --- |
| **I** | -20 ms to -5 ms | Silence | Silence | Latency period between stimulus and muscle response |
| **II** | -5 ms to 5 ms | Small positive, then large negative, and finally medium positive deflection | First field component in positive y- and z-direction; then small dip followed by a second smaller peak; and then flipping of the direction of both (y and z) field components | Propagating muscular action potential |
| **III** | 5 ms to 15 ms | Silence | Silence | Pre-activation period |
| **IV** | 15 ms 80 ms | Silence | Strong signal in the negative Y direction | (unknown presumably first interaction myosin-actin structures) |
| **V** | 80 ms to 300 ms | Silence | Magnetic activity in the y- and z-direction | Muscle contraction |

In order to obtain an approximation of when the contraction of the muscle starts, M-mode ultrasound scans have been performed before the OPM measurement. At the bottom of Figure 2D, an M-mode image is shown. This data suggests that the interval V corresponds to the muscle contraction. The source of the magnetic field during interval IV remains unknown.

### 3.1 Reproducibility and signal-to-noise ratio

In order to test the reproducibility of the MMG signals, 12 trials were referenced for the peak of the maximum positive signal in EMG 1 and are plotted together in Figure 3 for OPM 1 and 2 in the y- and z-direction and x- and z-direction (see Figure 3).

**Figure 3.**
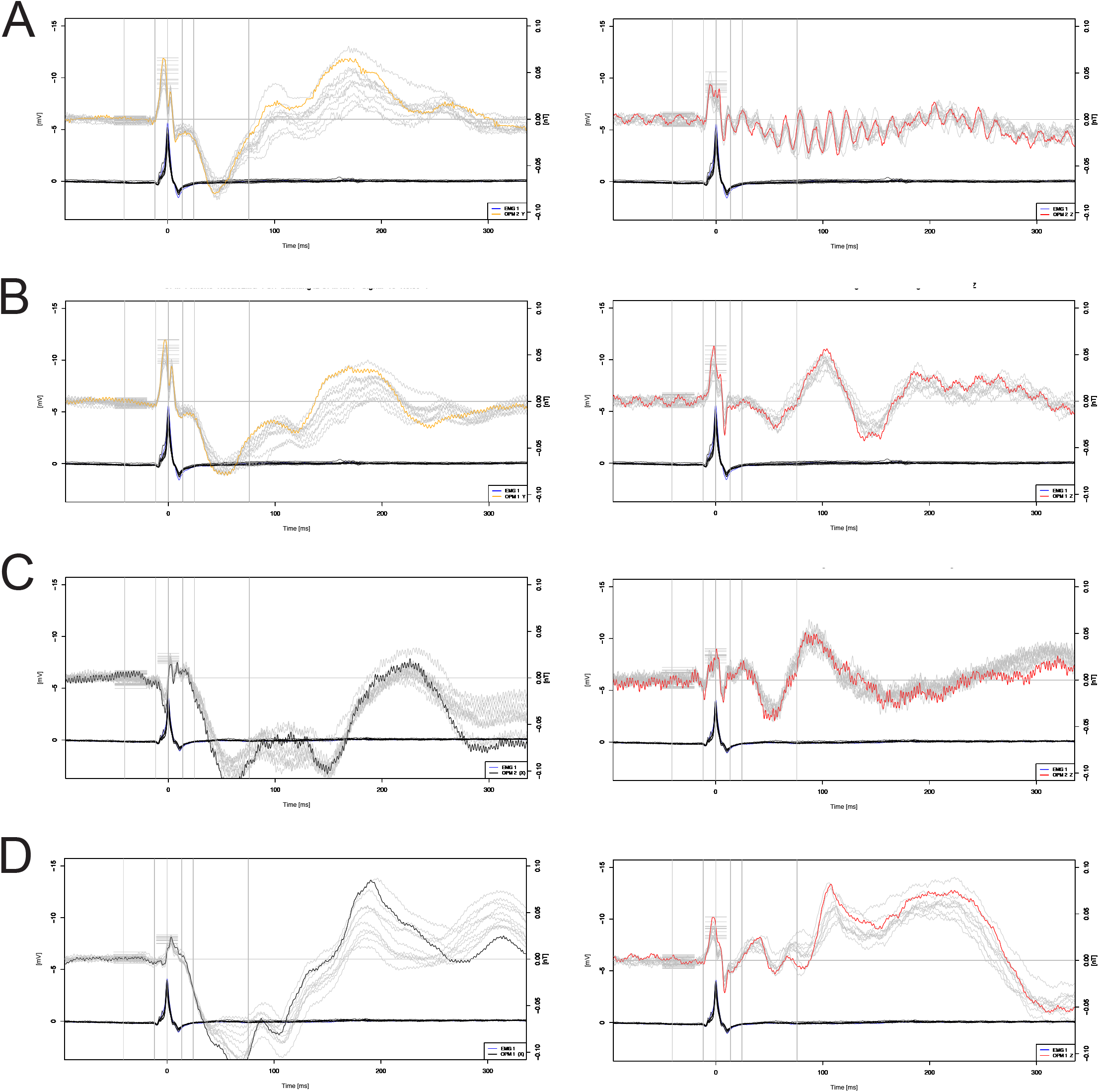
The reproducibility of the OPM signal was tested by time referencing 12 trials and plotting them in a single plot. This is depicted for sensors OPM 2 and 1 in the Y and Z direction (Panels A and B) and in the X and Z direction (Panels C and D). The small grey horizontal lines at −50 to −20 ms show the period during which the background signal was estimated. The small horizontal lines at the time of the peak show the time period during which the signal amplitude was estimated. The trial used for later analysis is shown colored. Remark: The z-direction is shown with an inverted signal to optimize visibility.

The signal amplitude of the background MMG signal was determined by measuring the difference of the maximum and minimum signal in the time period from −50 to −20 ms prior to the peak in EMG 1. The signal amplitude of the background was defined as the absolute difference between the minimum and maximum value. The foreground signal amplitude of the MMG signal was measured as the highest positive signal in the time period from −5 ms to 5 ms. The signal-to-noise ratio was calculated by dividing the foreground amplitude by the background amplitude.

Table 1 shows the signal-to-noise ratio of the MMG signal in the different directions and OPM sensor positions. For all sensors, the signal-to-noise ratio is above 3, and the magnetic field in the y-direction has the highest signal-to-noise ratio out of all sensors. The main variability in the ratio values is due to the varying intensity of the background signals. Specifically, the background signal is lowest in the y-direction (see Figure 3) and highest in the x-direction.

**Table 1.** Signal-to-noise ratio. (Median [Max – Min], n = 12)

| OPM Nr. | X-direction | Y-direction | Z-direction |
| --- | --- | --- | --- |
| 1 | 8.07 (16.78 - 2.16) | 9.78 (13.62 - 3.85) | 3.32 (4.66 - 1.54) |
| 2 | 3.95 (5.86 - 2.12) | 9.40 (31.19 - 3.37) | 3.15 (4.65 - 1.89) |

Given that the sensors had to be turned around the longitudinal axis in order to record the x direction, it was necessary to test if the sensors picked up the same signal prior to and after rotation. To compare the signal profiles, Figure 3 shows in Panels A and B (right side) the z-direction prior to the rotation and in Panel C and D (right side) the signal in the z-direction after axial rotation. If one compares Panel A and C with B and D, one can recognize that the signal during the period of the muscular action potential is the same before and after the rotation of the sensors. However, the later components show more variability and were therefore not further analyzed in this study.

Given the high reproducibility of the signal of the propagating muscular action potential and the excellent signal-to-noise ratio with values above three, further analysis was performed on individual trials without any averaging.

### 3.2 Correlation of MMG signal and EMG signal

In order to analyze the temporal characteristics of the EMG and MMG signals, the signals are plotted in Figure 4 with focus on the time period of −50 to 50 ms—the time period of the propagating muscular action potential—for each OPM sensor and all three directions (x, y, z). In addition to the OPM signal, each plot depicts the calculated bipolar EMG signal of the neighboring fine-wire EMG electrodes (i.e., d[EMG1, EMG2] for the bipolar EMG signal, which was calculated by V_EMG1_ - V_EMG2_). Both signals—the bipolar EMG signal and the MMG signal—have the same temporal onset and same duration. The MMG signals are strongest in the y direction, while the activity in the z direction is again larger than the activity in the x direction. Taking the finite wire model described in Broser (2020) into consideration, this suggests that the principal propagation direction of the muscular action potentials is in the x direction. When taking the simplified relationship between the scalar magnetic field strength B, the current I, the length of the wire (extend of the muscular action potential) s, and the distance between the sensor and current source (Westgard, 1997):

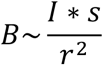

into consideration and estimates the current I by

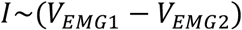

(with *V_PNQR_*, the voltage, measured at EMG Position 1), one can obtain the simplified relationship:

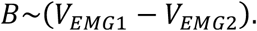

**Figure 4.**
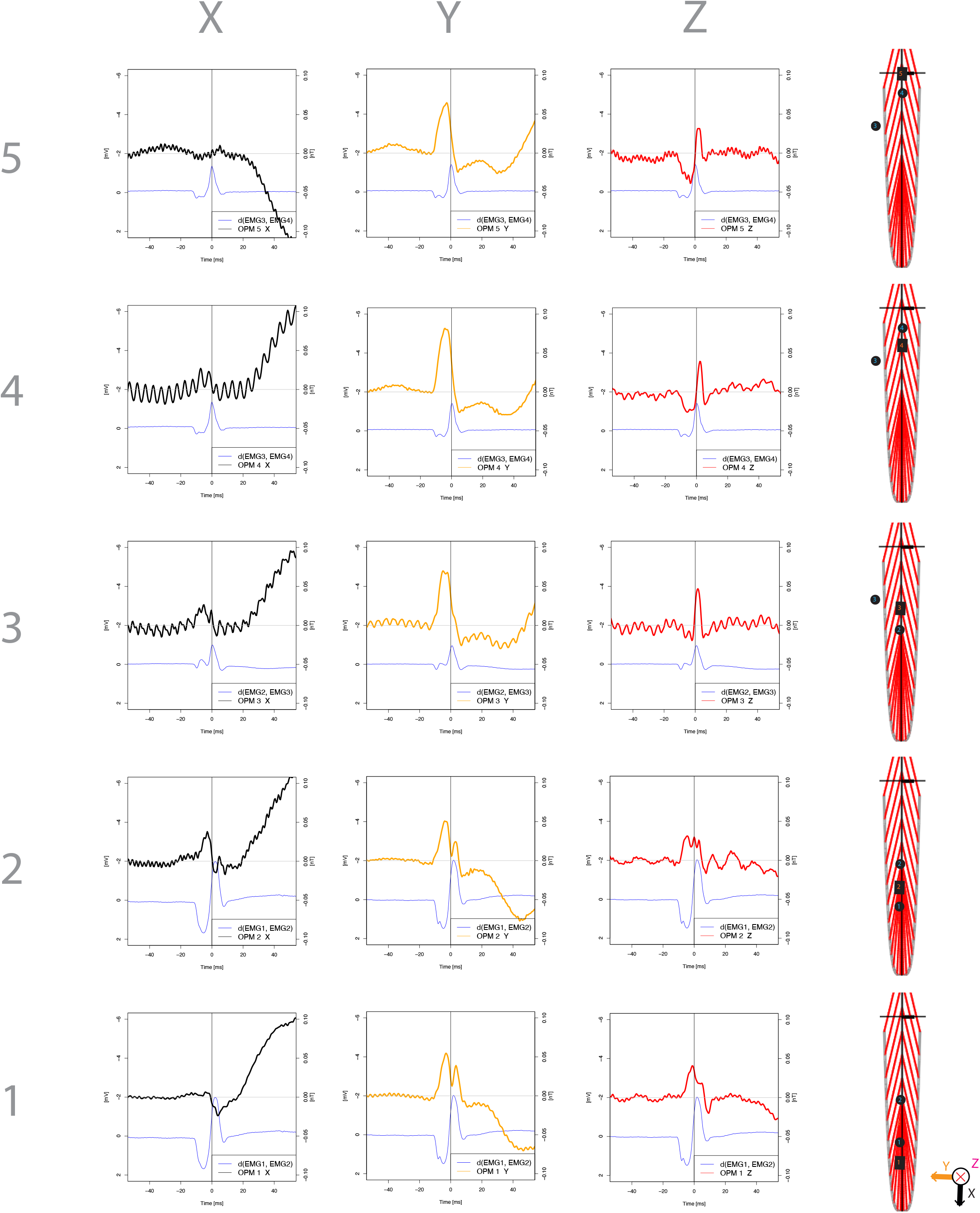
For each OPM sensor (1–5) the signal in the z-, y-, and z-direction in the time period of −50 ms to 50 ms around the muscular action potential is shown. In addition, the location of the sensor in relation to the muscle anatomy is depicted on the right side of the figure.

This linear relationship is identifiable in most of the MMG and EMG traces (see supplementary Figure 2). Given that the bipolar fine-wire EMG closely correlates with the current flowing in the tissue between the two electrodes as well as the fact that the magnetic field is generated by the current and not by the transmembrane membrane voltage, this finding is expected. Even so, this finding emphasizes the important fact that the MMG signal correlates with the electric current flowing in the muscle (in contrast, all other electrophysiology techniques measure the transmembrane voltage).

To further analyze the OPM signal profile, it is important to consider the precise anatomical structure of the muscle at the measurement position. The orientation of the muscular fiber, which determines the direction of the propagation of the muscular action potential, especially needs to be considered. Therefore, Figure 4 (right side) shows a model of the rectus femoris muscle, highlighting the fiber direction in the xy plane as well as the position of the selected OPM sensors and the neighboring fine-wire EMG positions.

### 3.3 Modelling the OPM Signal

Based on the magnetic field vector components, the direction of the propagating muscle action potential can be estimated. Due to the indirect stimulation by triggering a reflex response, the neuro muscular units are only partly in temporal synchrony. Therefore, both the EMG and the MMG signals are generated by muscular action potentials in different temporal phases. However, at the time of the signal onset, only one unit is activated, and, therefore, the direction can be estimated by the first deflection from the baseline. In order to do so, a parameter fitting approach as described in Broser (2020) and outlined in the method section was conducted. Figure 5 shows the estimated directions of the propagating muscular action potential at the different positions of the muscle. As expected, the direction vector turns from position OPM 4—about 20 degrees angulated—to facing almost straight forward for position OPM 1. Interestingly, for the position of OPM 5, we found a propagation direction parallel to the longitudinal extend of the muscle when we expected an angulation of about 20 degrees. In addition, the estimation of the direction for position OPM 3 is difficult given that the fist deflection in the z-direction is very small.

**Figure 5.**
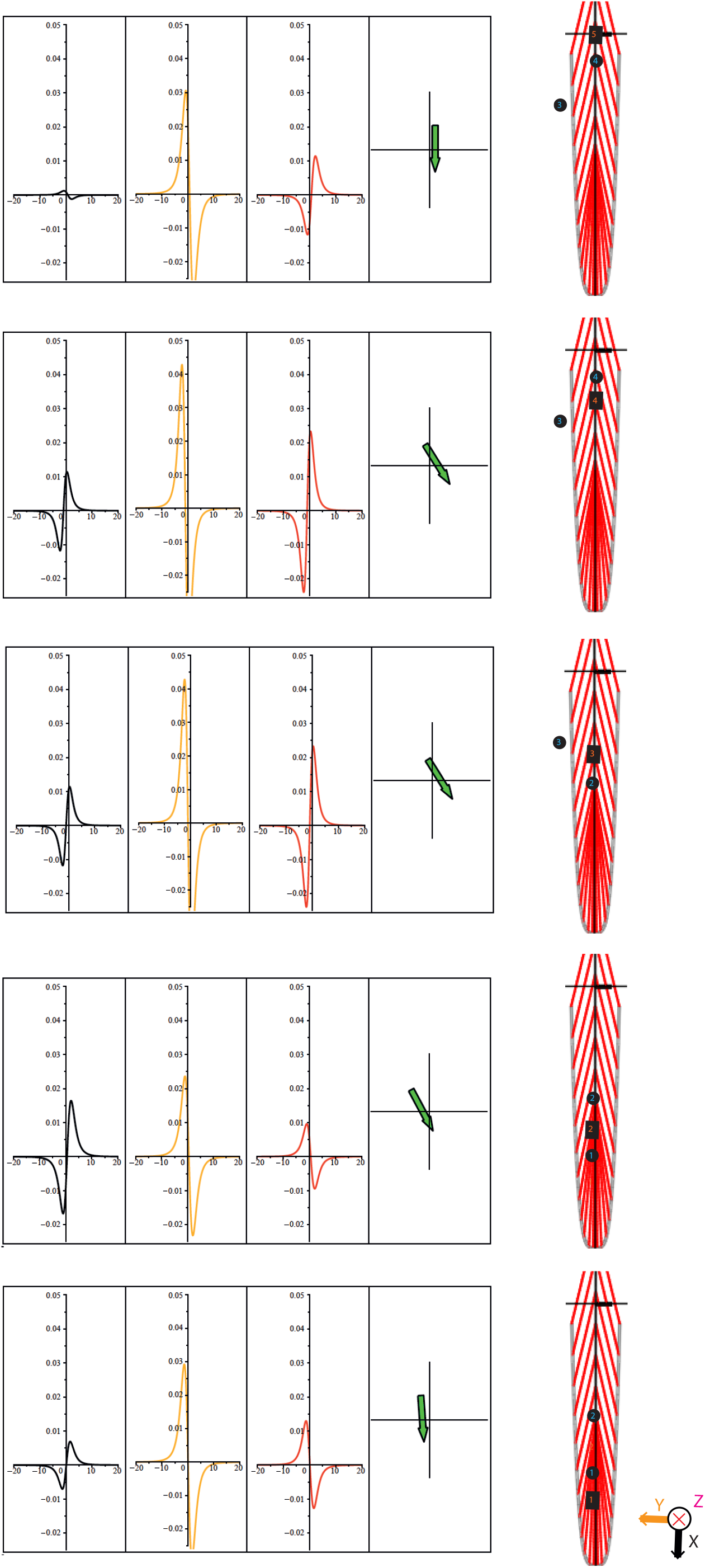
For each sensor, the signal onset in the x-,y-, and z-direction was used to model the magnetic field according to the finite wire model by Broser (2020). The resulting plots are shown in the middle; in addition, the direction of the simulated muscular action potential is shown on the right.

## 4 Discussion

In this study, we used optically pumped magnetometers to record the magnetic field components of the propagating muscular action potential along the fibers of the rectus femoris muscle. The muscle was activated by triggering a monosynaptic reflex response. About 20 ms after the hammer touched the skin of the subject, we could record a signal from the OPM devices and, at the same time, the fine-wire needle EMGs, which was similar to the latencies reported in the literature (Gürbüz, 2015). We could show that, after the muscular action potential, a latency period of about 10 ms follows. Then, a strong magnetic signal of unknown origin present in all three spatial directions can be recorded. We assume that this signal is due to the activity in the contractile apparatus or generated by movement. However, this remains to be clarified.

To test whether the stimulation of triggering a reflex results in a stable and reproducible response, we time-locked the MMG data to the peak of the EMG 1 electrode, plotted the data, and measured the signal-to-noise ratio, finding that this type of stimulation leads to a stable response with a high signal-to-noise ratio. To further analyze the muscular action potential, the analysis focused on the time period of −50 ms to 50 ms—around the peak of EMG 1. First, it was shown that a close temporal correlation between the EMG signal and the OPM signal exists. Previously, it was theoretically shown (Broser, 2020) that the MMG signal is generated by the current flow inside the muscular fiber. To obtain a gross estimate of the current flowing inside the rectus femoris muscle, we calculated the bipolar fine-wire EMG between the individual OPMs. Especially for OPMs 1 and 2, where the fibers are almost parallel to the longitudinal extend, the signal amplitude and curvature are in good correspondence. This result demonstrates that the magnetic field is, as theoretically expected, a result of the current flowing in the muscle, which is a completely new perspective of muscular physiology when compared to the techniques available so far, i.e., needle EMG measures the transmembrane voltage (Moritani, 2004).

The stimulation of the muscle by triggering a monosynaptic reflex response only leads to a weakly synchronized activity in the muscle. The neuro muscular units fire at slightly different timepoints. Therefore, the signal from muscle fibers with different time phases are recorded, making precise modelling difficult. However, at the very onset of the signals, modelling is possible; so, the direction of the muscular action potential can be estimated. Using this approach, we could reproduce the different pinnation angles inside the rectus femoris muscle given in the literature (Blemker, 2006). This finding demonstrates nicely how MMG can be used to analyze the direction of the muscle fibers.

While it was quite successful, this study had some important limitations. The OPM devices used have a restricted bandwidth, and, therefore, fine and fast components of the muscular action potential could not be recorded. In addition, the sensors used could only record the magnetic field in two directions, and, thus, to measure the signal in the third dimension, the sensors had to be manually turned during the experiment. Moreover, the anatomy of the rectus femoris is complex and changes significantly in short spatial intervals; therefore, a high-density grid of OPM sensors would be necessary to map the MMG precisely.

In conclusion, recording the magnetic field of the muscle not only allows for obtaining a non-invasive alternative for needle EMG recordings but also delivers information thus far inaccessible. First, the sensors record the current inside the muscle instead of the transmembrane voltage, and they can therefore reflect the spreading of excitation more precisely. Second, the vectorial character of the magnetic field discloses the direction of the current flow and therefore allows for drawing conclusions about the geometry of the fiber structure.

In addition to the magnetic components of the muscular action potential, we found additional magnetic field components about 20 ms after the action potential - the time we also see movement in the M mode ultrasound images - which we think originate from the contractile apparatus or from movement inside the muscle. If this is confirmed by further experiments, it could open up a whole new perspective of muscle physiology. For instance, this information would make it possible to link the propagating action potential with the activity of the contractile apparatus. From classical experiments, it is known that changes in the delay of the muscular action potential and the initiation of the interaction between myosin and action is closely correlated to the training level of the muscle (El-Ashker, 2019).

In summary we could show in this study that the magnetic field correlates with the current flowing in the longitudinal direction of the muscle fiber. The method is capable of showing the fiber direction in the muscle and possible allows the measurement of the electro mechanical delay.

So, MMG could open a new analysis method for the contractile apparatus in athletes as well as in patients suffering from neuromuscular diseases. A first prove of a principal study (currently in preparation) shows that pathological spontaneous muscular activity can be recorded by MMG (Marquetand et al., in preparation).

In order to extend our knowledge regarding the sources of late magnetic field components and to optimize the modelling of the muscular action potential, we plan to conduct a series of animal experiments on isolated muscle fibers. These experiments will help to specifically understand the late magnetic field, which we currently assume to be generated by the contractile apparatus.

## Abbreviations

MMG: magneto myography
OPM: optically pumped magnetometer
MAP: muscle action potential

## Acknowledgement

We want to thank Jürgen Dax for the construction of the non-magnetic reflex hammer and the support during the measurements.

---

**Supplementary Figure 1.**
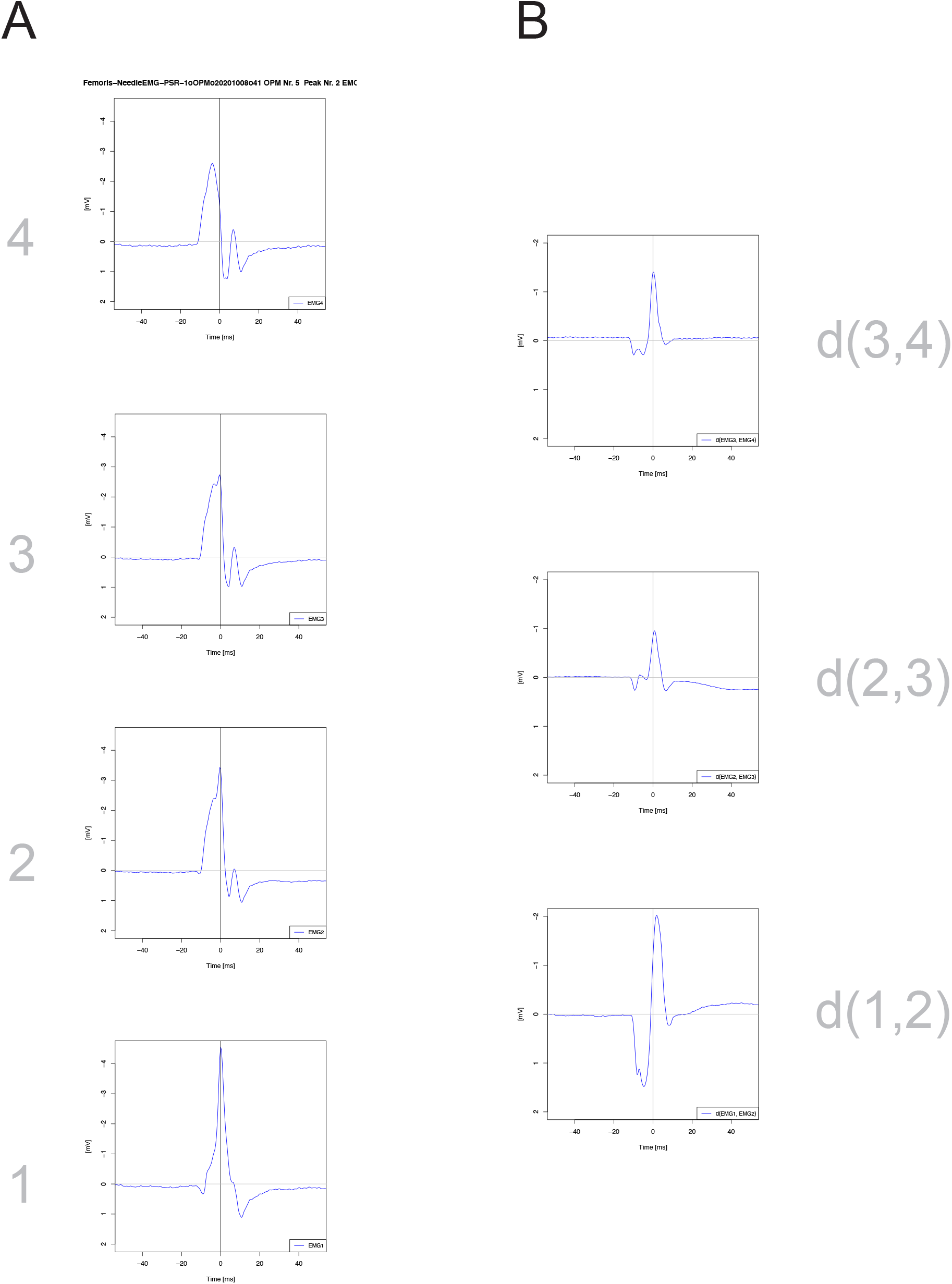
This figure shows the monopolar EMG signal for each fine-wire electrode on the left and the bipolar EMG signal on the right.

**Supplementary Figure 2.**
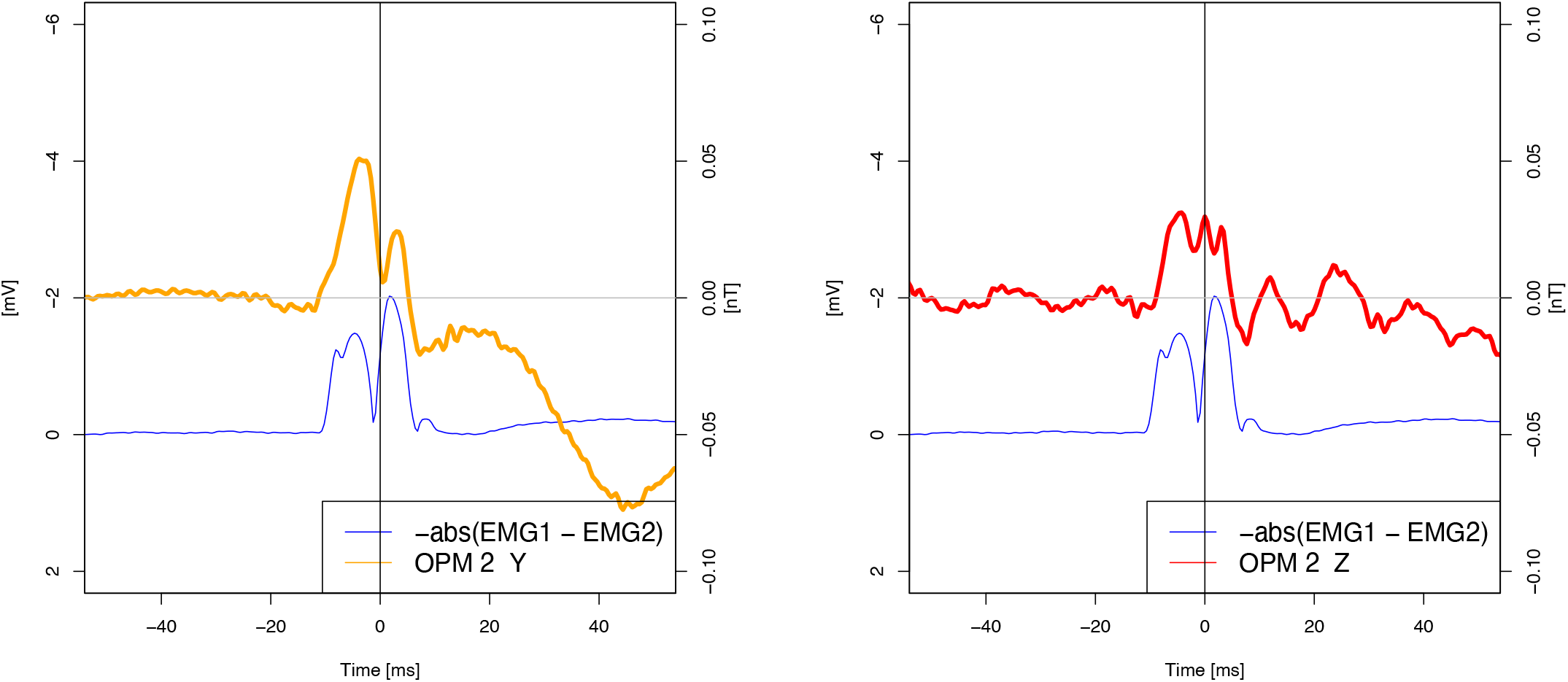
OPM sensor 2 is between EMG 1 and 2. Assuming that the current flows in a straight line along the X axis, the magnetic field strength should show a linear correlation with the bipolar EMG signal (only the magnitude of the EMG signal is shown).

## Footnotes

1 Bipolar EMG needles are ferromagnetic and therefore cannot be used with OPM devices.

